## Supplementary Figures 1-7 for "TMEM16 and OSCA/TMEM63 proteins share a conserved potential to permeate ions and phospholipids"

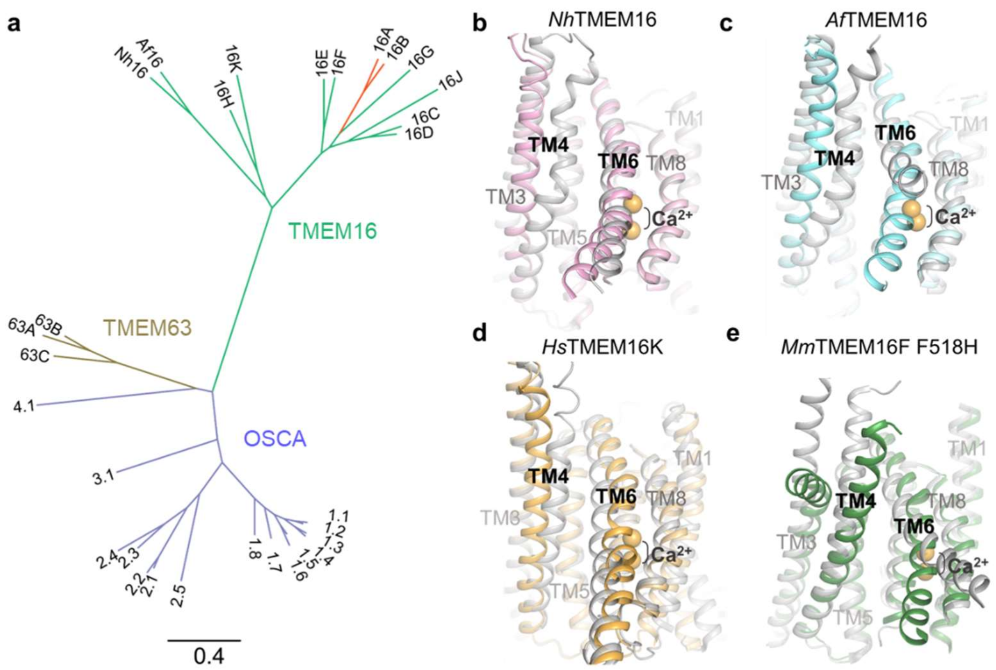


f

```
                 TM 4
MmTMEM16F    QMATSITASIISFIIIMILNTIYEKVAIMITNFE 539
MmTMEM16A    RVTVTATAVIINLVVIILLDEVYGCIARWLTKIE 564
HsTMEM16K    SVLLY-VPSIIYAIVIEIMNRLYRYAAEFLTSWE 388
AfTMEM16     GYLVF-IPTILVSALIPTMSAVLLTVATKLNDYE 350
NhTMEM16     QYLGF-LPTIFLVIGTPTISGVLMGAAEKLNAME 358
AtOSCA1.2    SVIQGFLPGIALKLFLAFLP----SILMIMSKFE 452
MmTMEM63A    PVISQFFPTLLLWSFSALLP----SIVYYSTLLE 489
HsTMEM63B    PIITQFFPTLLLWCFSALLP----TIVYYSAFFE 502
MmTMC1       K----NEMNMVMSLLGMFCP----TLFDLFAELE 525
                                             *
```

**Supplementary Figure 1: Conformational changes in transmembrane helices (TMs) 4 and 6 are associated with TMEM16 scramblase activities.** (**a**) Phylogenetic tree of TMEM16 (CaCC, red; putative and validated CaPLSase, green), OSCA (purple), TMEM63 (gold) families. Structure alignments of (**b**) fungal *Nh*TMEM16 in calcium-bound open (PDB 6QM9, pink) and calcium-bound closed (PDB 6QMB, gray), (**c**) fungal *Af*TMEM16 in calcium-bound open (PDB 6E0H, cyan) and calcium-free closed (PDB 6DZ7, gray), (**d**) human TMEM16K in calcium-bound open (PDB 5OC9, orange) and calcium-bound closed (PDB 6R7X, gray), and (**e**) mouse TMEM16F F518H in calcium-bound open (PDB 8B8J, green) and calcium-free closed (PDB 8B8G, gray) conformations. (**f**) Sequence alignment of TM 4 region for mouse TMEM16F, mouse TMEM16A, human TMEM16K, fungal *Af*TMEM16, fungal *Nh*TMEM16, thale cress OSCA1.2, mouse TMEM63A, human TMEM63B, and mouse TMC1. Yellow and cyan highlighting denotes residues identified in this work and hydrophobic gate residues previously identified, respectively.

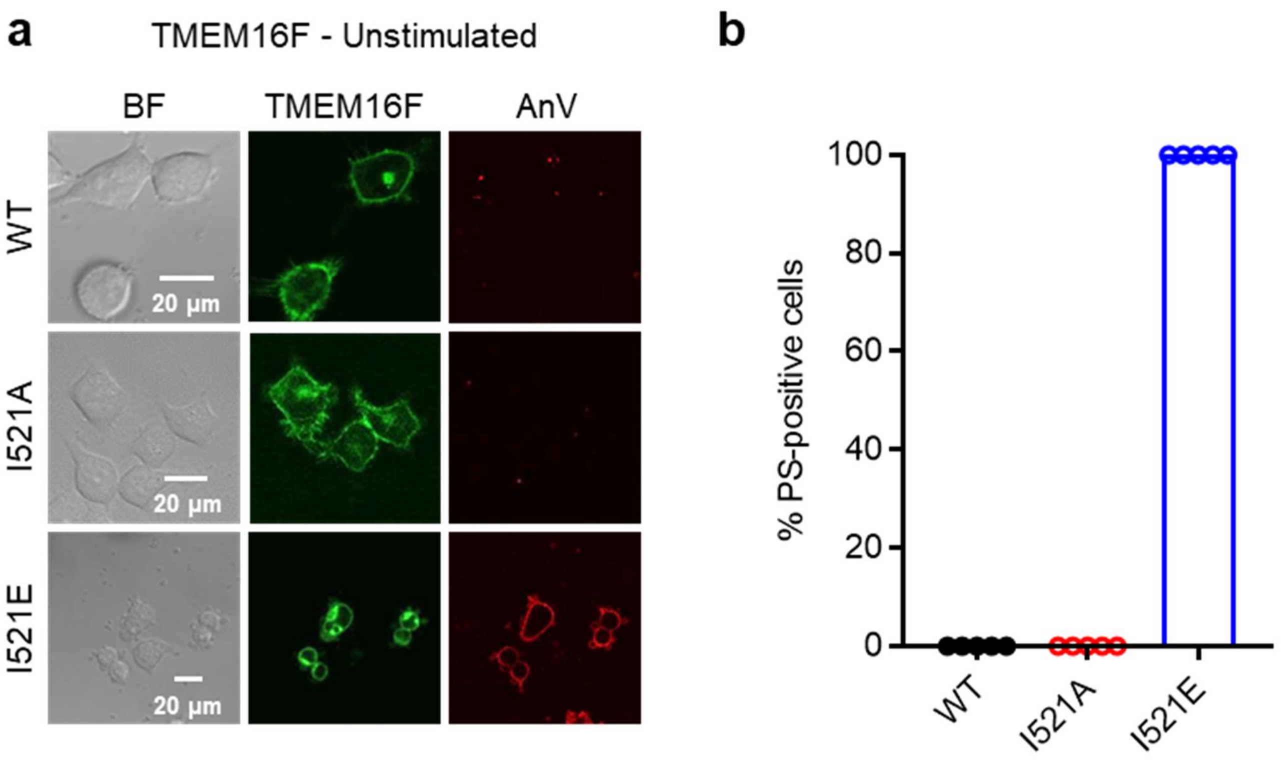


**Supplementary Figure 2: Alternative side chains at I521 cause differential PS exposure.** (**a**) Representative images of TMEM16F knockout (KO) HEK293T cells expressing eGFP-tagged TMEM16F wildtype (WT), I521A, and I521E (center column). CF 594-conjugated Annexin V (AnV, right column) labelled PS exposing cells. BF denotes bright field images (left column). (**b**) Quantification of the percentage of cells with AnV labelling for TMEM16F WT (n=5), I521A (n=5), and I521E (n=5) transfected cells. Values were derived from images of biological replicates. Statistical comparisons were not conducted due to zero variance within each group.

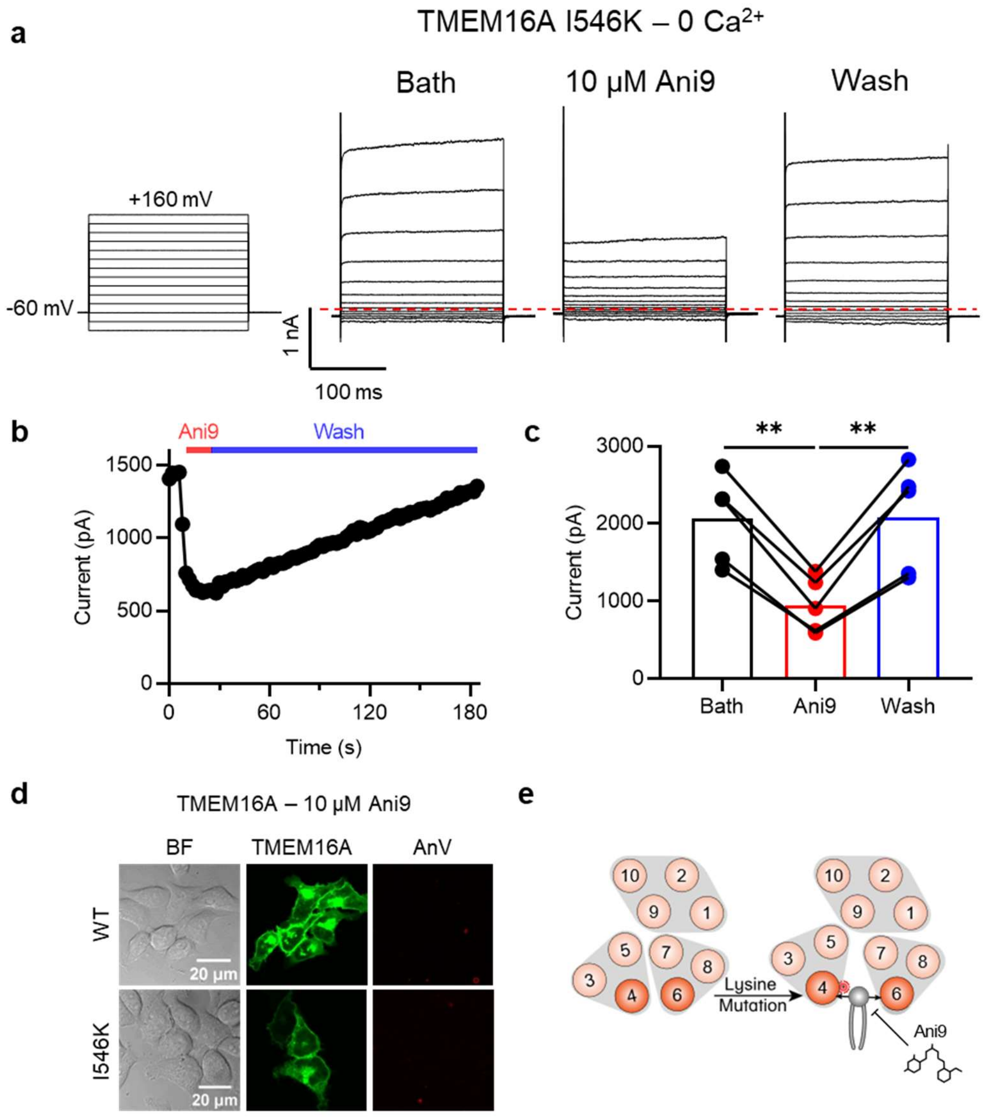


**Supplementary Figure 3: Ani9 attenuates ion channel and phospholipid scramblase activities of TMEM16A I546K.** (**a**) Representative whole-cell current recordings elicited by the voltage protocol shown (left) for TMEM16A I546K before, during, and after application of 10 μM Ani9. (**b**) Representative time-course of Ani9 application and wash-off. (**c**) Quantification of peak current from (**b**) before (bath), during (Ani9), and after (wash) Ani9 application (n=5). (**d**) Representative images of TMEM16F KO HEK293T cells expressing eGFP-tagged TMEM16A WT or I546K (center columns). CF 594-conjugated AnV (right columns) failed to label any PS exposing cells (n=4). BF denotes bright field images (left columns). (**e**) Ani9 attenuates mutant-induced phospholipid permeability.

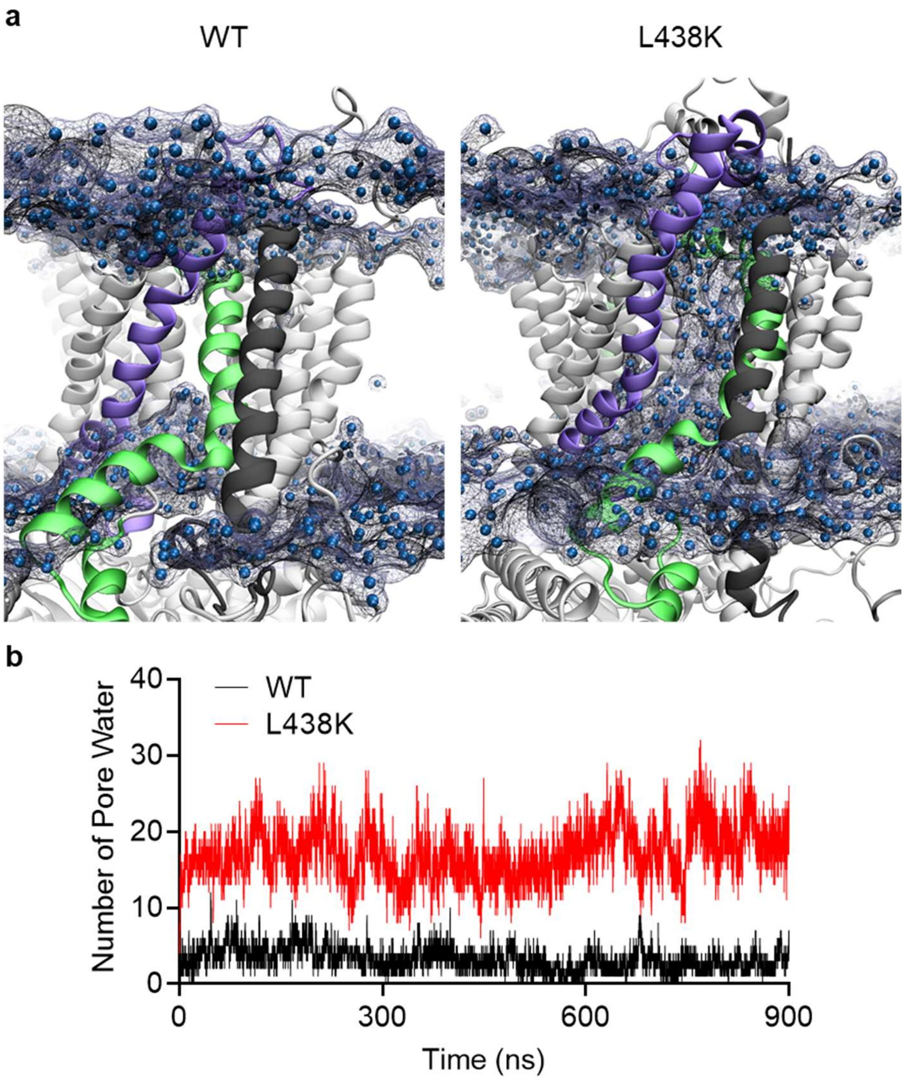


**Supplementary Figure 4: Atomistic MD simulations of OSCA1.2 WT and L438K exhibit differential hydration of the pore region.** (**a**) Representative timepoints (900 ns) for OSCA1.2 WT (left) and L438K (right). (**b**) Quantification of water occupancy in the pore region for WT (black) and L438K (red) over the duration of the simulation. The pore region was defined as within 7 Å of I432, which is centrally positioned in the upper pore.

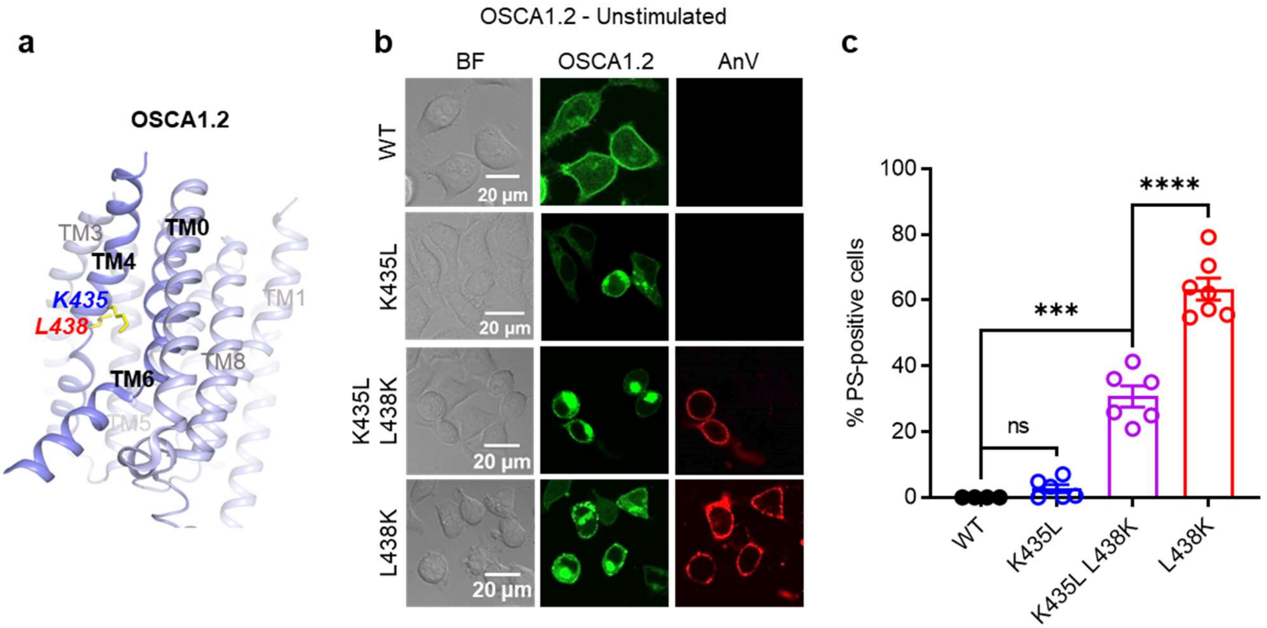


**Supplementary Figure 5: Role of K435 in OSCA1.2 mutant phospholipid permeability.** (**a**) TM 4 mutant locations mapped onto the TM 4/6 interface of OSCA1.2 (PDB 6MGV) with key residues shown as yellow sticks. (**b**) Representative images of TMEM16F KO HEK293T cells expressing eGFP-tagged OSCA1.2 WT, K435L, K435L/L438K, or L438K mutants (center column). CF 594-conjugated AnV (right column) labelled PS exposing cells. BF denotes bright field images (left column). (**c**) Quantification of the percentage of cells with AnV labelling for OSCA1.2 WT (n=4), K435L (n=6), K435L/L438K (n=6), and L438K-transfected cells (n=7). Statistical comparisons were conducted with unpaired t-tests with Welch's correction (ns: p>0.05, ***: p<0.001, ****: p<0.0001).

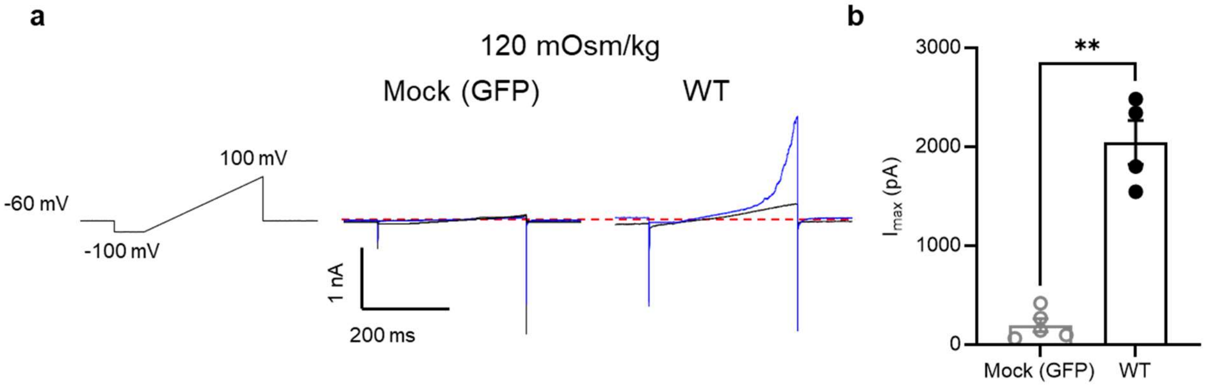


**Supplementary Figure 6: Hypotonic stimulation activates the OSCA1.2 ion channel.** (**a**) Representative current recordings elicited by the voltage protocol shown for mock (GFP) and OSCA1.2 WT-transfected cells before (black) and after (blue) reduction of bath osmolarity from 310 to 120 mOsm/kg. (**b**) Quantification of peak currents from (**a**) for mock (n=5) and WT (n=4) after reduction of bath osmolarity. Statistical comparison was conducted with an unpaired t-test with Welch's correction (**: p<0.01).

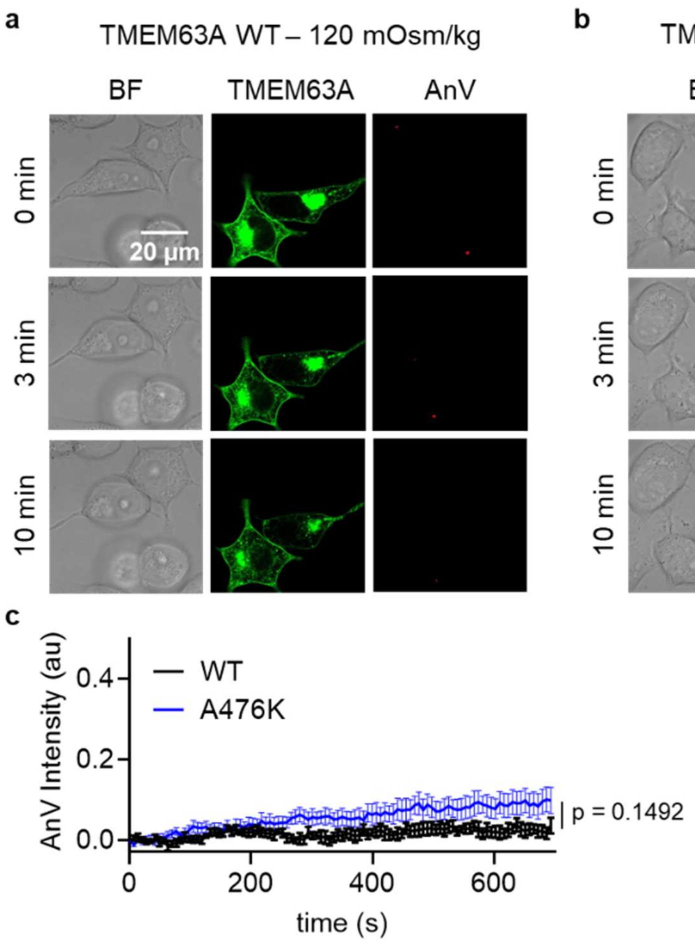


**Supplementary Figure 7:** TMEM63A hypotonic stimulation. (**a-b**) Representative images of hypotonic osmolarity stimulation of TMEM16F KO HEK293T cells expressing eGFP-tagged TMEM63A (**a**) WT or (**b**) the A476K mutant (middle columns). CF 594-conjugated AnV (right columns) labelled PS exposing cells. BF denotes bright field images (left columns). Each row of representative images corresponds to the indicated time after hypo-osmotic stimulation. (**c**) Quantification of AnV intensity for TMEM63A WT (n=3) and A476K (n=5) after hypo-osmotic stimulation. Statistical comparison was conducted with an unpaired t-test with Welch's correction.
